# A Genetic Atlas of Direct and Inverse Neuropsychiatric-Cancer Comorbidity

**DOI:** 10.64898/2026.07.10.737193

**Authors:** María Flores-Rodero, Jaume Forés Martos, Joan Vicent Sánchez Ortí, Salvador Martínez, Frank Winkler, Alfonso Valencia, Rafael Tabarés-Seisdedos, Jon Sánchez-Valle

**Affiliations:** Department of Medicine, University of Valencia, Valencia, Spain; Computational Biology Group, Life Sciences Department, Barcelona Supercomputing Center, Barcelona, Spain; TMAP – Evaluation Unit in Personal Autonomy, Dependency and Serious Mental Disorders, University of Valencia, Valencia, Spain; Department of Pathology, Stanford University School of Medicine, Stanford, CA, USA; Catalan Institution for Research and Advanced Studies, Barcelona, Spain; INCLIVA Health Research Institute, Valencia, Spain; Center for Biomedical Research in Mental Health Network (CIBERSAM), Health Institute Carlos III, Madrid, Spain; Instituto de Neurociencias de Alicante, Universidad Miguel Hernández–Spanish National Research Council (UMH-CSIC), San Juan de Alicante, Spain; Neurology Clinic and European Center for Neurooncology (EZN), Heidelberg University Medical Faculty, University Hospital Heidelberg, Heidelberg, Germany; Clinical Cooperation Unit Neurooncology, German Cancer Consortium (DKTK), German Cancer Research Center (DKFZ), Heidelberg, Germany

**Keywords:** genetic correlation, GWAS, pleiotropy, inverse comorbidity, cancer neuroscience, neuropsychiatric disorders

## Abstract

Direct and inverse comorbidities between neuropsychiatric disorders and cancer are increasingly recognised as important features of the nervous system–cancer relationship, yet the inherited genetic architecture underlying these patterns remains poorly understood. Here, we analysed pairwise genetic correlations across 35 diseases represented by 115 GWAS datasets, including 9 psychiatric disorders, 10 neurological diseases and 16 cancers, using linkage disequilibrium score regression (LDSC) and high-definition likelihood (HDL), complemented by meta-analysis, subtype-resolved analyses, local covariance mapping and multi-omic benchmarking. Genetic correlations were predominantly positive and strongest within disease categories, whereas cancer–neurological pairs showed the weakest overall genetic affinity. Meta-analysis and subtype resolution uncovered associations obscured in aggregate analyses, including opposing correlations between familial and late-onset Alzheimer’s disease and lung cancer, revealing subtype-dependent neuro-oncological biology. Local analyses identified recurrent genomic loci where direct comorbidities are consistent with shared inflammatory, interferon, survival and tissue-remodelling programs, whereas inverse comorbidities suggest competing demands on apoptotic regulation, immune tone and stress-response calibration between neuronal and tumour-cell states. Together, these findings provide a genome-scale genetic framework for neuropsychiatric–cancer comorbidity and identify shared inherited biological programs as candidates for mechanistic investigation and therapeutic translation.

## 1 Introduction

The nervous system and cancer engage in bidirectional interactions that influence tumour growth, immune dynamics and disease progression, giving rise to the field of cancer neuroscience^1–3^. A defining feature of these interactions is that neuropsychiatric and neurological disorders exhibit markedly non-random patterns of co-occurrence with cancer: some conditions co-occur more frequently than expected, whereas others are mutually exclusive^4–8^. For instance, individuals with schizophrenia show a lower prevalence of lung cancer after adjustment for smoking^9^, genes upregulated in Alzheimer’s disease are frequently downregulated in lung cancer^10^, and antidepressant use has been linked to a reduced risk of colorectal cancer^11^. The protective effect of some neuropsychiatric conditions against cancer appears to extend to first-degree relatives, suggesting that inherited factors contribute to these relationships^12, 13^. The asymmetry between direct and inverse neuropsychiatric–cancer comorbidity has therefore motivated growing interest in the inherited biological programs that may link the nervous system and cancer, yet the genome-wide genetic architecture underlying these patterns remains poorly characterised.

The prevalence of comorbidities is influenced by sex, age, ancestry and pharmacological treatment^11, 14^, and their burden is substantial: psychiatric disorders and neoplasms together account for nearly 40% of the global chronic disease burden, with mental and neurological disorders contributing 10.4% of all disability-adjusted life years and cancer 7.6%^15^. Individuals with mental illness show an elevated risk of physical comorbidities overall (OR = 1.84), including cancer, and dementia frequently co-occurs with psychosis and schizophrenia^16–18^. Understanding the biological basis of these patterns is therefore both scientifically and clinically urgent.

Research on the molecular basis of disease co-occurrence has revealed extensive pleiotropy across the human genome, with 81% of studied genes associated with at least two traits^19^. When altered, these pleiotropic genes influence biological drivers involved in the pathogenesis of multiple disease categories^6, 8^. Transcriptomic analyses have proven particularly useful for generating biological hypotheses for disease co-occurrence^20, 21^. Large-scale studies using UK Biobank data have provided valuable insights into the genetic basis of multimorbidity^22^, yet most analyses focus on co-occurring conditions and overlook inverse comorbidities and their mechanistic potential. Genome-wide genetic correlation approaches offer a complementary strategy by capturing shared inherited architecture upstream of downstream molecular states and before disease onset, treatment exposure or tissue degeneration distort the signal^23, 24^.

Here, we provide a genome-scale atlas of direct and inverse comorbidity across neuropsychiatric and cancer phenotypes. We analysed 35 unique diseases represented by 115 GWAS summary-statistic datasets and computed genome-wide genetic correlations using LDSC and HDL, detecting both positive and negative correlations across psychiatric, neurological and cancer trait pairs. To identify biological processes contributing to these relationships, we applied local covariance analysis with SUPERGNOVA, followed by pathway enrichment with rGREAT, and benchmarked the findings against 10 independent omics layers. Results are compiled into an interactive Shiny app for open exploration.

## 2 Results

### 2.1 Landscape of genetic correlations across neuropsychiatric and cancer traits

To map the shared genetic architecture between neuropsychiatric and cancer phenotypes, with disease classifications assigned according to predominant symptomatology (Supplementary Note 1), we analysed genetic correlations across 35 unique diseases: 9 psychiatric, 10 neurological or neurodegenerative and 16 cancer phenotypes, represented by 115 GWAS summary-statistic datasets. We used two complementary methods, LDSC and HDL (Methods and Supplementary Note 1). LDSC computed correlations for all 6,555 disease pairs, whereas HDL computed 5,873 correlations; 682 pairs were unavailable owing to low heritability or overlapping populations (Supplementary Note 2). Across the 5,873 trait pairs computed by both methods, LDSC and HDL showed high concordance (Pearson’s *r* = 0.93; Supplementary Fig. 1a), with 86% of pairs directionally consistent. When restricting to pairs significant in both methods, only four pairs presented discordant directionality, and all involved the same minor bipolar disorder dataset (BIP-PGC-2011), with HDL estimates collapsing to values effectively rounded to zero owing to reference-panel mismatch rather than genuine method disagreement. Overall, HDL identified a higher fraction of significant correlations than LDSC (odds ratio [OR] = 1.66, *P* = 2.45 *×* 10^−41^), particularly for negative correlations (OR = 2.57, *P* = 7.92 *×* 10*^−^*^13^) and inter-category pairs (OR = 2.09, *P* = 4.81 *×* 10*^−^*^38^).

Genetic correlations were predominantly positive (approximately 73–76%; Supplementary Tables 1 and 2) and markedly enriched within disease categories. Intra-category pairs were significantly more likely to yield significant correlations than inter-category pairs (LDSC: OR = 6.47, *P* = 4.69 *×* 10*^−^*^235^; HDL: OR = 4.82, *P* = 7.22 *×* 10*^−^*^174^). Among category pairs, psychiatric–psychiatric pairs showed the strongest affinity (LDSC: OR = 58.65, *P ≈* 0; HDL: OR = 29.13, *P ≈* 0), whereas cancer–neurological pairs showed the weakest genetic overlap (LDSC: OR = 0.17, *P* = 7.73 *×* 10*^−^*^66^; HDL: OR = 0.21, *P* = 1.72 *×* 10*^−^*^62^). Full Fisher exact test results by software are provided in Supplementary Table 3, and software comparisons are provided in Supplementary Table 4. Agreement between methods, quantified by the Jaccard index, was highest within psychiatric–psychiatric pairs (0.94) and lowest for cancer–neurological pairs (0.43; Supplementary Fig. 2). Jaccard breakdowns by category pair are shown in Table 1.

**Table 1:** Jaccard index of concordance between LDSC and HDL by disease-category pair. Values are shown separately for global, positive and negative significant correlations.

| Category pair | Global | Positive | Negative |
| --- | --- | --- | --- |
| PSY-PSY | 0.94 | 0.94 | 0.00 |
| CAN-CAN | 0.56 | 0.56 | 0.35 |
| NEU-NEU | 0.57 | 0.57 | 0.50 |
| CAN-PSY | 0.53 | 0.56 | 0.31 |
| NEU-PSY | 0.56 | 0.61 | 0.31 |
| CAN-NEU | 0.43 | 0.40 | 0.54 |

Among notable neuro-oncological relationships (Fig. 1), ADHD showed strong positive associations with lung cancer (LDSC *r_g_*= 0.356; HDL *r_g_*= 0.397) and cervical cancer (LDSC *r_g_* = 0.354; HDL *r_g_* = 0.348), consistent with reported genetic links between neurodevelopmental disorders and these cancer types, as well as higher smoking rates in ADHD^25–27^. Autism spectrum disorder showed a negative correlation with prostate cancer (LDSC *r_g_* = *−*0.176; HDL *r_g_* = *−*0.146), replicating an inverse comorbidity previously proposed through transcriptomic analysis^28^. Among neurological traits, multiple sclerosis was positively associated with renal cancer (LDSC *r_g_* = 0.287) and anxiety (LDSC *r_g_* = 0.383), matching the established psychiatric comorbidity profile of multiple sclerosis^29, 30^. Schizophrenia showed a negative correlation with melanoma (LDSC *r_g_* = *−*0.118), consistent with earlier reports of inverse cancer associations in psychotic disorders^31^. Alzheimer’s disease showed a single consistent positive correlation with ALS (LDSC *r_g_* = 0.242; HDL *r_g_* = 0.304), supporting reported genetic and clinical overlap^32, 33^. No significant associations with cancer were detected for Alzheimer’s disease in the main-dataset analyses, motivating the subtype-resolved and meta-analytic approaches described below.

**Fig. 1:**
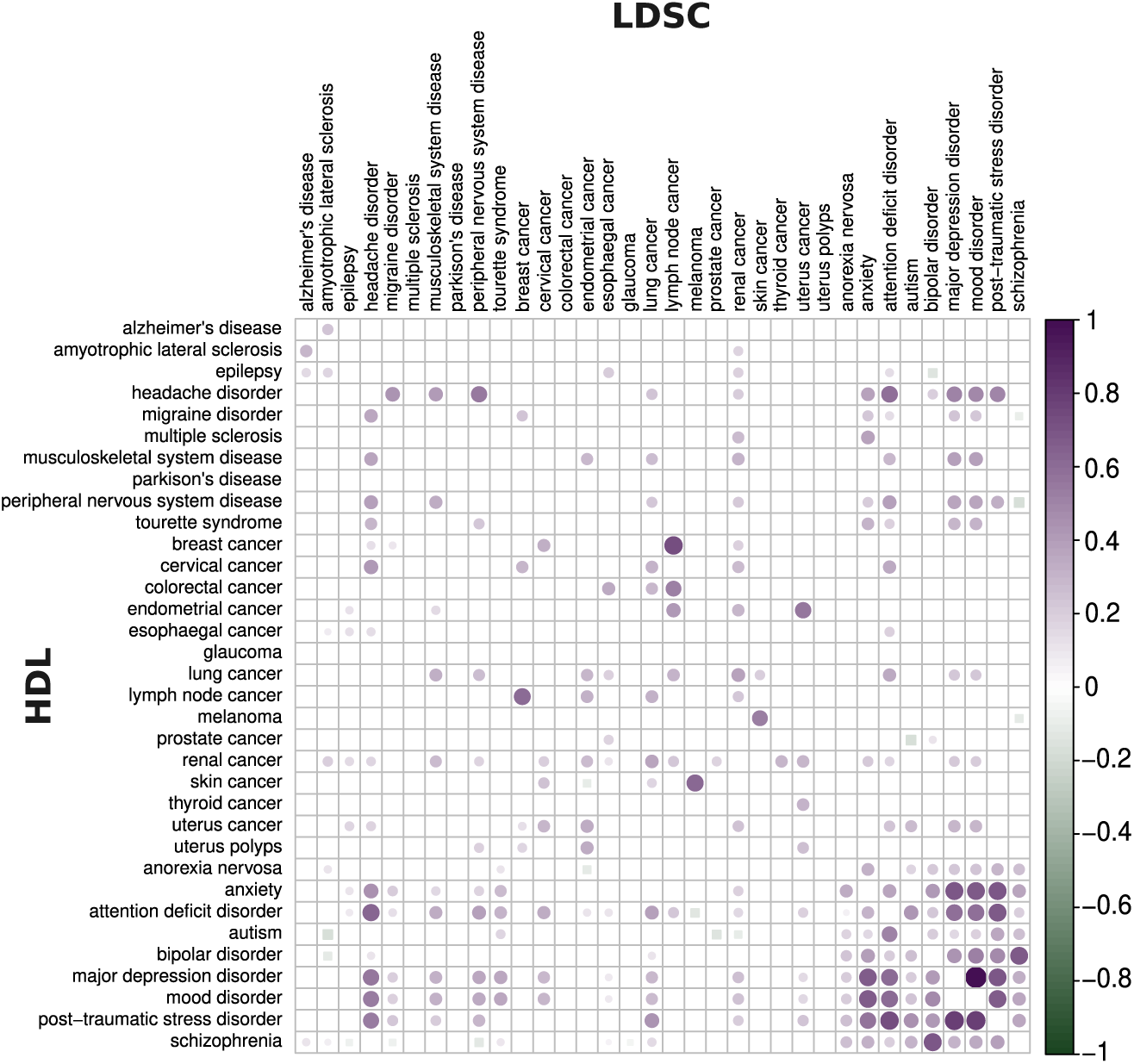
Global genetic correlation heatmap estimated by LDSC and HDL. Pairwise genetic correlations (*r_g_*) estimated using LDSC (upper triangle) and HDL (lower triangle). Only FDR-significant correlations among main datasets are displayed. Purple circles denote positive correlations and green squares denote negative correlations.

### 2.2 Meta-analysis reveals subtype-sensitive neuro-oncological signals

To increase sensitivity beyond single-dataset analyses, we meta-analysed correlation estimates across datasets representing the same disease pairs. The meta-analysis showed a concordance of 0.96 with the original results for both methods (Supplementary Fig. 3) and recovered additional significant neuro-oncological associations not visible in the main-dataset scan (Fig. 2).

**Fig. 2:**
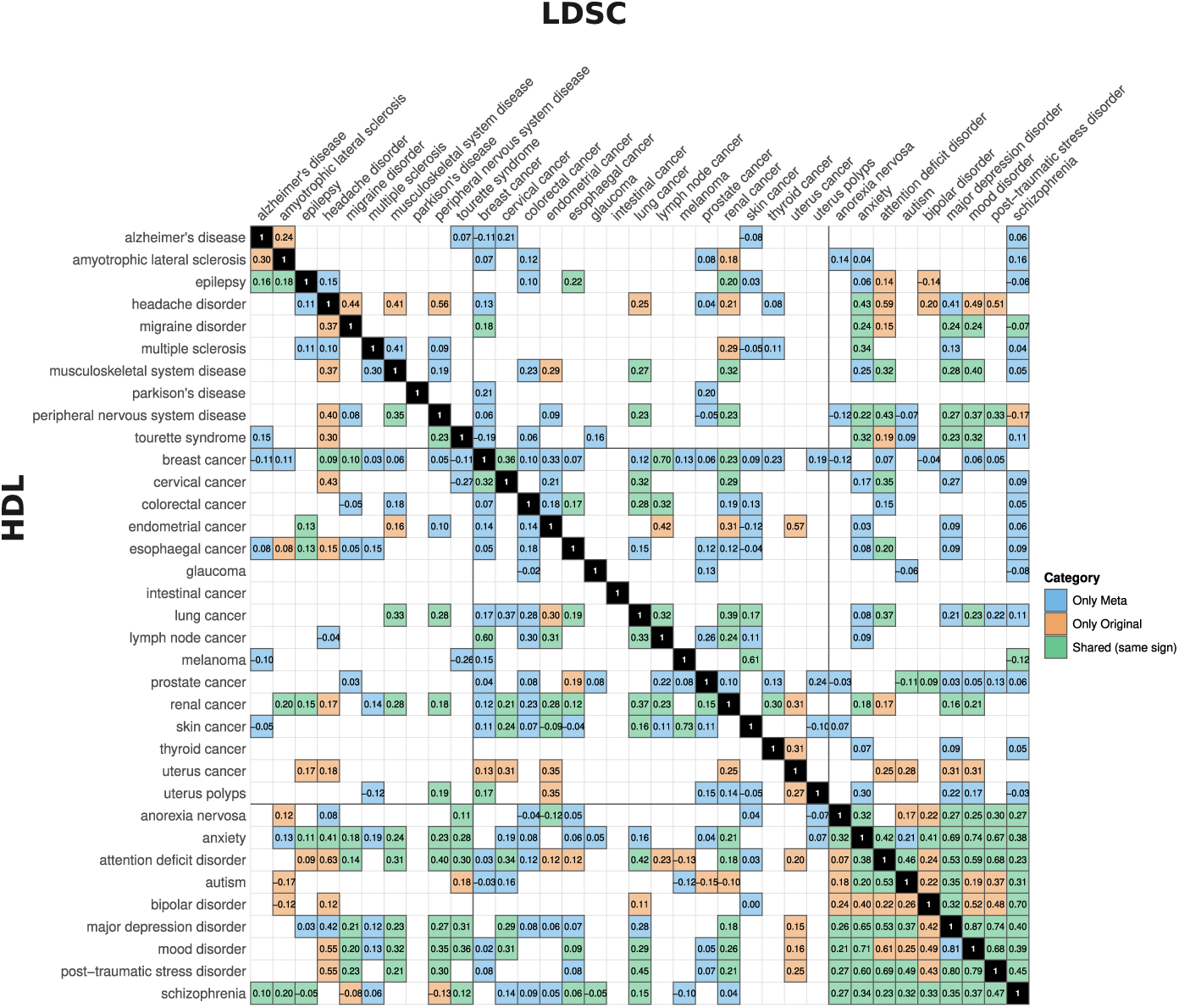
Meta-analysis genetic correlation heatmap. Meta-analysis results of genetic correlation profiles estimated by LDSC (upper triangle) and HDL (lower triangle) across main and minor datasets. Cell colour indicates agreement with the original analyses: blue marks correlations detected only in the meta-analysis, orange marks correlations unique to the original data and green marks correlations shared between both, in which case the value shown corresponds to the original estimate.

For Alzheimer’s disease, meta-analysis identified negative correlations with breast cancer in both methods (LDSC *r_g_* = *−*0.107; HDL *r_g_* = *−*0.105) and a negative correlation with melanoma in HDL (*r_g_* = *−*0.103), consistent with previous epidemiological and Mendelian randomisation studies^34–36^. A positive correlation with cervical cancer was also detected in LDSC (*r_g_*= 0.206). For Parkinson’s disease, which showed no significant signal in the main-dataset analysis, meta-analysis revealed positive correlations with breast cancer (LDSC *r_g_* = 0.210) and prostate cancer (LDSC *r_g_* = 0.198), although the direction of the Parkinson’s disease–prostate cancer relationship remains contested across epidemiological and meta-analytic studies^37–40^. For ALS, aggregation uncovered additional positive correlations with breast cancer (LDSC *r_g_* = 0.070; HDL *r_g_* = 0.106), colorectal cancer (LDSC *r_g_* = 0.123) and schizophrenia (LDSC *r_g_* = 0.160). For multiple sclerosis, meta-analysis revealed a strong positive correlation with musculoskeletal disease in both methods (LDSC *r_g_* = 0.411; HDL *r_g_* = 0.303), compatible with recognised comorbidity burden^30, 41^, as well as additional associations with thyroid cancer in LDSC (*r_g_* = 0.111) and oesophageal cancer in HDL (*r_g_* = 0.145).

### 2.3 Subtype and sex-stratified analyses uncover divergent neuro-oncological biology

Further significant associations emerged when considering subtype and minor datasets. The most striking finding from subtype analyses was that familial Alzheimer’s disease and late-onset Alzheimer’s disease showed opposite directions of genetic correlation with both lung cancer and ADHD, a pattern with direct implications for how genetic comorbidity studies should handle clinically heterogeneous disease categories. Familial Alzheimer’s disease showed strong negative correlations with lung cancer (LDSC *r_g_* = *−*0.372; HDL *r_g_* = *−*0.462), with similar effect sizes in family-history datasets including maternal history (LDSC *r_g_* = *−*0.523; HDL *r_g_* = *−*0.477), and additional negative correlations with renal cancer (LDSC *r_g_*= *−*0.316; HDL *r_g_*= *−*0.244) and squamous-cell skin cancer (LDSC *r_g_* = *−*0.338; HDL *r_g_* = *−*0.324). Late-onset Alzheimer’s disease showed positive correlations with ADHD datasets in HDL (*r_g_*= 0.137), in contrast to familial Alzheimer’s disease datasets (LDSC *r_g_* = *−*0.376; HDL *r_g_* = *−*0.244). This divergence is biologically coherent: familial Alzheimer’s disease is dominated by penetrant amyloid-processing biology involving APP, PSEN1 and PSEN2, whereas late-onset Alzheimer’s disease is shaped more strongly by APOE, lipid handling, vascular ageing and immune regulation^42, 43^. Genetic comorbidity studies that aggregate Alzheimer’s disease subtypes without distinction are therefore likely to obscure or cancel opposing neuro-oncological signals, and the same caveat applies to any disease for which clinically or aetiologically distinct subtypes have been included under a single phenotype label (Fig. 3).

**Fig. 3:**
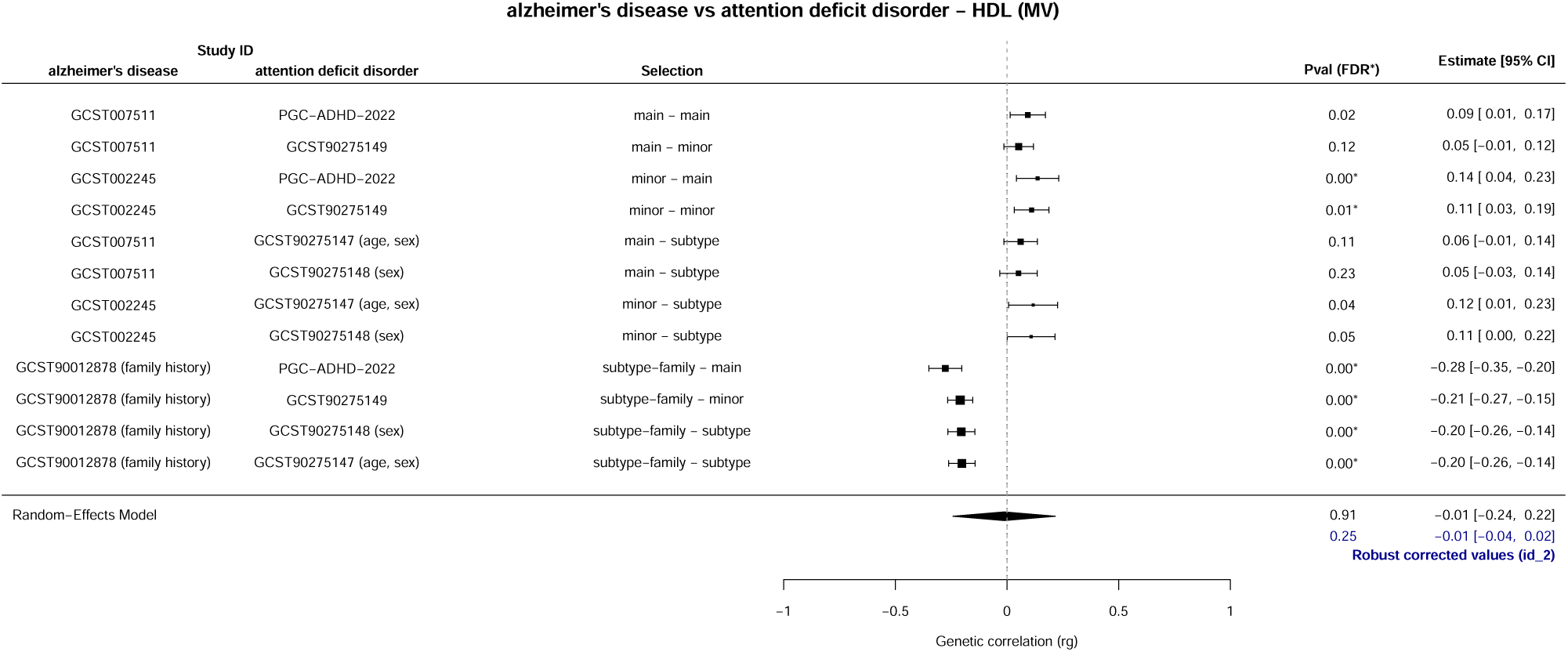
Forest plots of subtype- and stratified-analysis genetic correlations estimated by HDL. Forest plots summarise HDL *r_g_* estimates and 95% confidence intervals for the Alzheimer’s disease and ADHD comparisons discussed in the text, including analyses stratified by subtype, family history, sex and lung cancer subtype. Points represent estimated genetic correlations, horizontal lines show 95% confidence intervals and the vertical reference line indicates *r_g_* = 0.

Further subtype-dependent effects were observed for ADHD and lung cancer. Effect sizes were larger for squamous-cell carcinoma than for adenocarcinoma (LDSC *r_g_*= 0.391; HDL *r_g_*= 0.518 versus LDSC *r_g_*= 0.234; HDL *r_g_*= 0.345), suggesting that the neurodevelopmental–oncogenic genetic overlap is histologically specific. Sex-stratified analyses further indicated stronger schizophrenia–lung cancer associations in males than females: in the main lung cancer dataset, correlations were larger in males (HDL *r_g_* = 0.173) than females (HDL *r_g_* = 0.107), and for squamous-cell lung carcinoma, the association was significant in males (HDL *r_g_* = 0.200) but not in females. Squamous-cell skin carcinoma showed negative correlations with anorexia nervosa (LDSC *r_g_* = *−*0.119; HDL *r_g_* = *−*0.140) and bipolar disorder (LDSC *r_g_* = *−*0.133), and a negative correlation was observed between partial epilepsy and schizophrenia (LDSC *r_g_* = *−*0.151), with a stronger effect in males (LDSC *r_g_*= *−*0.168).

### 2.4 Recurrent genomic loci and biological programs underlying neuro-oncological comorbidity

To identify the genomic regions and biological programs underlying global correlations, we conducted local covariance analysis followed by pathway enrichment (Methods). We focus here on four representative disease pairs that illustrate the major neuro-oncological themes identified: Alzheimer’s disease–colorectal cancer, ADHD–lung cancer, familial Alzheimer’s disease–lung cancer and bipolar disorder–epilepsy. Results for additional pairs (ADHD–familial Alzheimer’s disease, schizophrenia–lung cancer, autism–prostate cancer and ADHD–melanoma) are provided in Supplementary Note 3.

#### Alzheimer’s disease–colorectal cancer

The positive local correlation between late-onset Alzheimer’s disease and colorectal cancer was supported by a single significant genomic region on chr5:140.4–143.4 Mb, enriched for FGFR ligand binding and downstream signalling pathways (FDR = 0.006), including isoform-specific FGFR1 and FGFR2 activation, SHC-and FRS-mediated cascades and oncogenic PI3K signalling. Key genes in this region include FGF1/FGFR1, which supports neuronal survival and is implicated in Alzheimer’s disease biology^44, 45^; NDFIP1, which regulates iron homeostasis and *β*-amyloid pathology in Alzheimer’s disease and acts as a tumour suppressor in cancer^46, 47^; and NR3C1, whose methylation has been associated with neuroinflammation, dysregulated amyloid-*β* metabolism and tumour progression^48, 49^. A negative region at chr22:45.4–45.8 Mb containing PHF21B^50, 51^ and NUP50^52^ was also identified, although no significant pathway enrichment survived FDR correction.

#### ADHD–lung cancer

ADHD showed significant positive correlations with multiple lung cancer subtypes, with consistent pathway enrichment across datasets. The most recurrent signal was telomere and chromosome maintenance, enriched across squamous carcinoma, lung carcinoma and smoker-stratified comparisons, driven by the chr9:126.9–128.9 Mb locus. This region contained LHX2, a key regulator of cortical development^53, 54^; PBX3, involved in malignant progression in lung cancer models^55^; and HSPA5/GRP78, a stress chaperone implicated in lung cancer progression^56, 57^. Interferon-*α/β* signalling and antiviral response pathways reached significance specifically in adenocarcinoma comparisons (FDR = 0.015–0.045), consistent with a type I interferon–JAK–STAT immune axis at this locus and with clinically actionable immune and survival pathways in non-small-cell lung cancer^58–60^. This interferon–JAK–STAT axis is of particular interest because sympathetic neural signalling can antagonise type I interferon responses in the tumour microenvironment, suggesting that the genetic architecture identified here may modulate the balance between pro- and anti-tumour neural inputs^61^. In family-history lung cancer subtypes, the enrichment profile shifted toward vesicle trafficking and cytoskeletal organisation (FDR = 0.006–0.013), suggesting that different ascertainment strategies capture partially distinct shared loci. A second recurrent region at chr2:103.2–104.5 Mb contained POU3F3, a neurodevelopmental gene whose pathogenic variants cause neurodevelopmental disorders and which has also been linked to proliferation, migration and invasion in cancer^62, 63^ (Fig. 4).

**Fig. 4:**
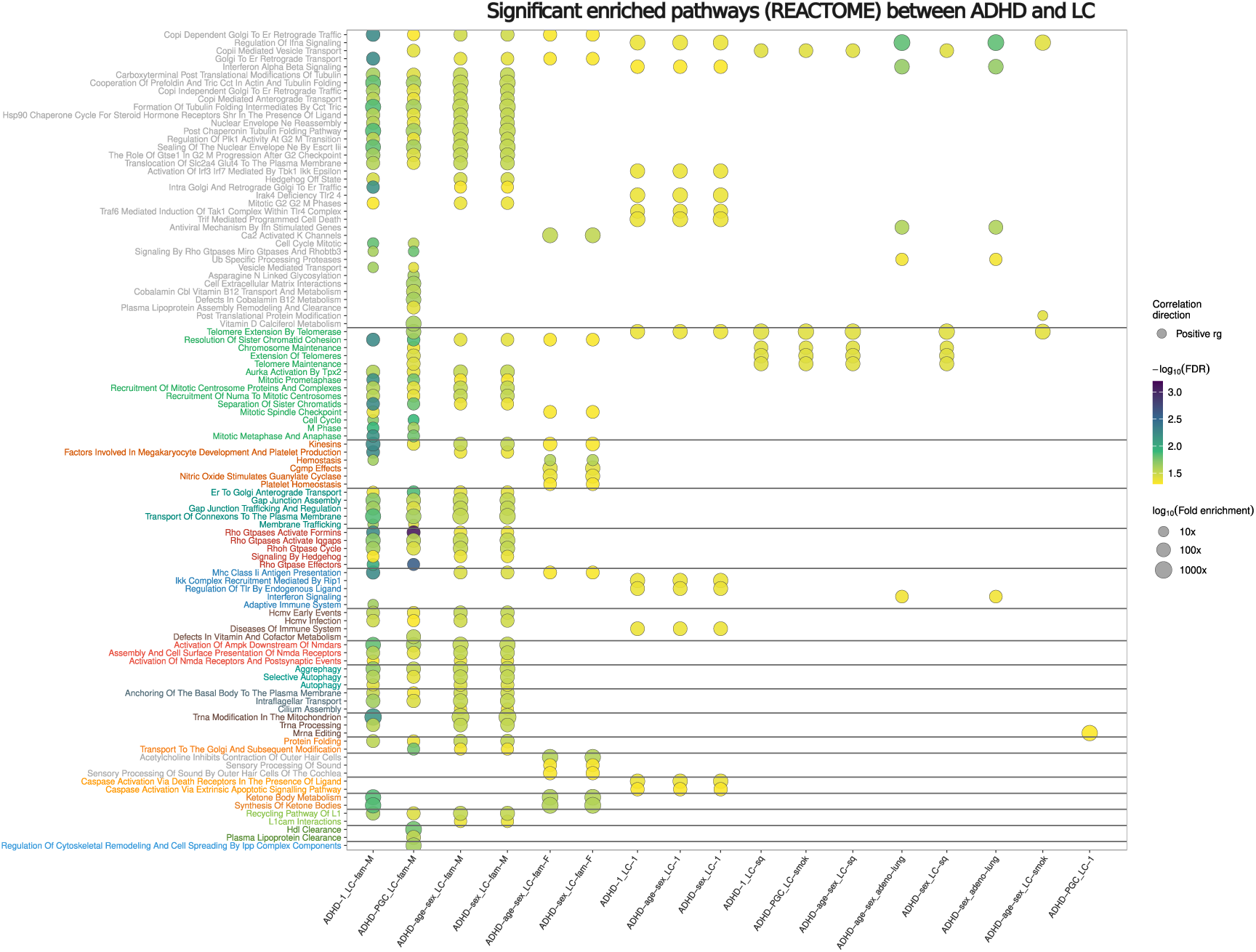
Reactome pathway enrichment across ADHD–lung cancer analyses. Reactome pathways enriched in significant local-correlation regions across the ADHD–lung cancer comparisons, highlighting recurrent telomere maintenance, interferon signalling, vesicle trafficking and cytoskeletal-organisation programs.

#### Familial Alzheimer’s disease–lung cancer

The correlation between familial Alzheimer’s disease and lung cancer was subtype-dependent and directionally mixed, exemplifying how the same genomic architecture can generate opposing neuro-oncological signals depending on disease ascertainment. Negative local signals on chr16:29.7–31.4 Mb and chr17:65.0–66.3 Mb were enriched for immune regulation and inflammatory signalling, with genes encoding components of the MAPK/ERK cascade^64, 65^ and transcriptional regulation by the RUNX3 cluster (FDR = 0.016). The familial-by-mother comparison showed mixed local effects: a positive signal at chr12:67.0–67.8 Mb enriched for AMPA receptor trafficking and iron uptake (FDR = 0.016), and a negative region at chr13:46.5–46.9 Mb enriched for the complement cascade and angiotensinogen metabolism (FDR = 0.005–0.013). In smoker-stratified analyses, rRNA processing and modification pathways were enriched (FDR = 0.007–0.016), pointing to translational stress and nucleolar biology as a shared neuro-oncological axis. A positive region at chr21:47.5–48.1 Mb in the smokers’ comparison was enriched for acetyl-CoA metabolism and protein acetylation, with genes including COL6A1/COL6A2, linked to neuroprotective responses against amyloid-*β* toxicity^66^, and S100B, an astrocytic factor that becomes pro-inflammatory at high levels and is overexpressed around amyloid-*β* plaques^67^ (Fig. 5).

**Fig. 5:**
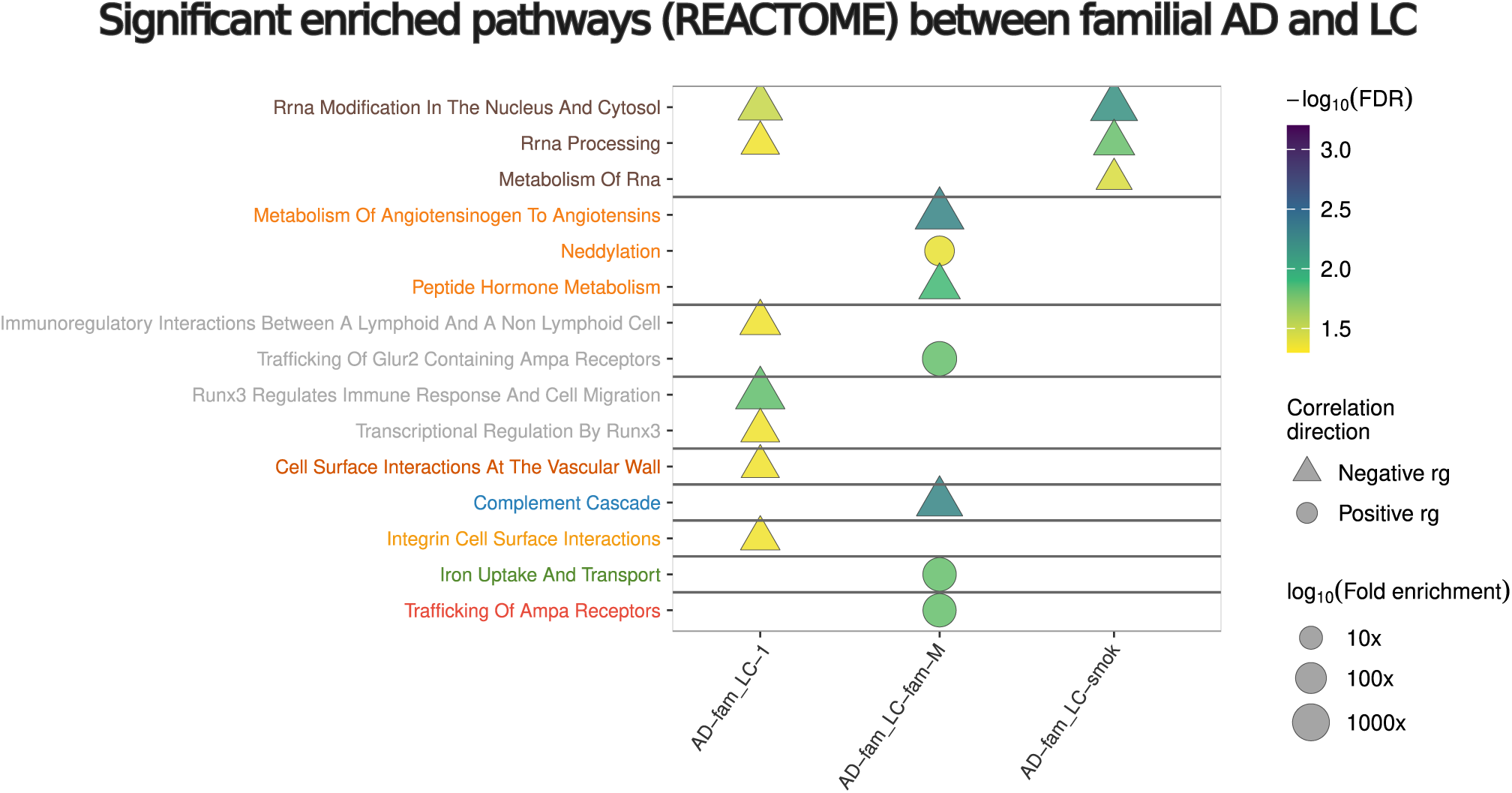
Reactome pathway enrichment across familial Alzheimer’s disease–lung cancer analyses. Reactome pathways enriched in significant local-correlation regions across the familial Alzheimer’s disease–lung cancer comparisons, highlighting recurrent programs related to RNA metabolism and processing, immune and inflammatory signalling, cell-surface and integrin-mediated interactions, hormone and angiotensin metabolism and AMPA-receptor trafficking.

#### Bipolar disorder–epilepsy

The global negative correlation between bipolar disorder and epilepsy was localised to bidirectional local signals. Negative components were driven by regions on chr3:52.2–54.0 Mb and chr12:2.17–2.88 Mb, enriched for membrane depolarisation and stress-response pathways, including HSF1 activation, the Hsp90 chaperone cycle and adrenergic signalling (FDR *<* 0.031). Key genes included CACNA1C^68^ and CACNA1D, where gain-of-function variants have been reported in autism and epilepsy^69^; PRKCD, linked to epileptogenesis through Fyn–PKC*δ* signalling^70^; and the ITIH1–ITIH3–ITIH4 cluster, a replicated bipolar susceptibility locus involved in extracellular matrix stabilisation and perineuronal biology^71, 72^. The shared positive locus at chr2:103.2–104.5 Mb, via POU3F3, links this pair to the ADHD–lung cancer comparison above, illustrating the transdiagnostic reach of individual genomic modules. The negative chr12 locus was also detected in schizophrenia–lung cancer associations in females (Supplementary Note 3.2).

### 2.5 Multi-omic convergence on epidemiologically established comorbidities

To place the genetic correlations in a broader biological context, we examined whether epidemiologically established comorbid pairs could be independently recovered across ten omic layers, including the LDSC and HDL results from this study (Fig. 6).

**Fig. 6:**
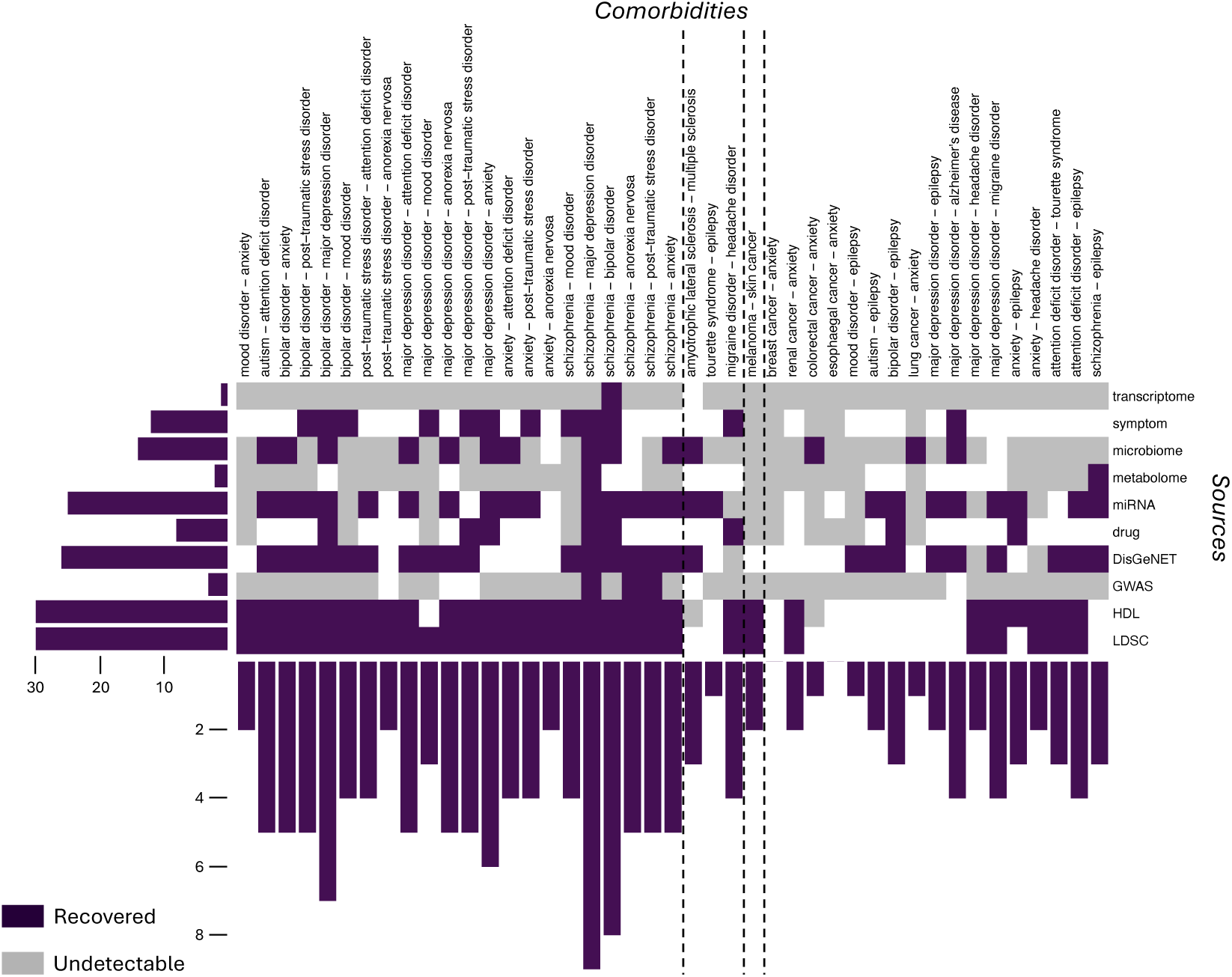
Multi-omic recovery of epidemiologically established comorbidities. Each column represents a disease pair identified as comorbid in Danish electronic health records. Rows correspond to ten evidence sources: genetic correlation methods from this study (LDSC and HDL), GWAS Catalog co-associations, gene–disease associations (DisGeNET), drug target overlap, miRNA, metabolome, microbiome, gut microbiome, symptom profiles and transcriptomic similarity. Purple indicates that the comorbidity is recovered by the given omic layer; white indicates no significant similarity detected; grey indicates that the pair is undetectable owing to missing data for at least one disease in that resource. Dashed vertical lines delineate disease-category pairs. The left bar plot shows the number of established comorbidities recovered from each omic source. The bottom bar plot shows the number of omic sources that independently recover each disease pair.

Genetic correlation methods ranked among the most productive sources of recovery across all analysed pairs. Transcriptome, microbiome and metabolome data showed more limited coverage, in part owing to higher rates of missing data for at least one disease in a pair. The pattern of recovery was markedly non-uniform across disease categories: psychiatric pairs showed the highest multi-omic convergence, with several pairs, including major depression–anxiety, bipolar disorder–schizophrenia and ADHD–PTSD, recovered by five or more independent sources. By contrast, inter-category pairs involving neurological and cancer diseases were recovered by fewer layers and showed a higher proportion of undetectable entries, consistent with the weaker and more heterogeneous genetic correlations reported above and suggesting that their shared architecture may be more tissue-dependent, stage-specific or restricted to particular subtypes.

At the level of individual omic sources, LDSC and HDL recovered as many or more established comorbidities than most other layers, supporting their utility as discovery tools for neuropsychiatric–cancer comorbidity research. The convergence between genetic and transcriptomic evidence is particularly notable: inherited susceptibility and downstream gene expression capture complementary aspects of disease biology, and their co-recovery of the same pairs strengthens the case for a shared molecular aetiology. Pairs recovered exclusively or predominantly by genetic correlation but not by other layers are of particular interest, as they may reflect comorbidities that are genetically determined but not yet visible at the molecular level in current databases, either because the relevant tissues or disease stages have not been adequately profiled, or because the shared architecture operates through regulatory rather than coding variation.

### 2.6 Summary of shared neuro-oncological genetic programs

Together, the findings described above converge on a coherent framework of shared and opposing genetic programs linking neuropsychiatric and cancer phenotypes. Figure 7 summarises the study design, the key disease-pair relationships identified and the biological programs underlying direct and inverse comorbidity. Direct comorbidities are consistent with shared inflammatory, survival, tissue-remodelling and telomere-maintenance programs that may serve adaptive functions in both neuronal and tumour contexts, whereas inverse comorbidities suggest opposing selective pressures on apoptotic regulation, stress-response proteostasis, neuroprotection and immune-tone calibration. The context-dependent balance of these programs, rather than any single pathway, may determine whether a given neuropsychiatric–cancer relationship manifests as co-occurrence or mutual exclusion.

**Fig. 7:**
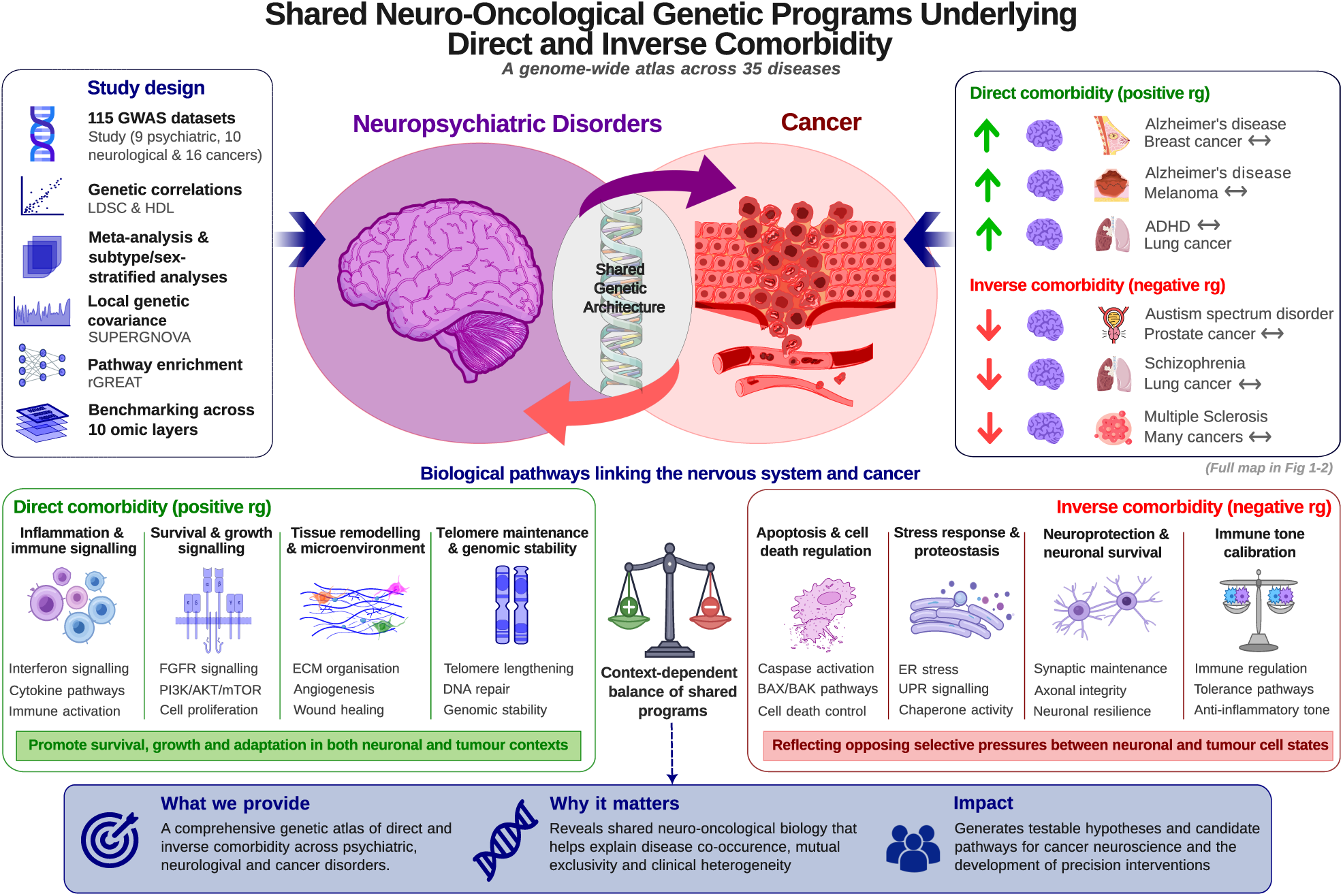
Shared neuro-oncological genetic programs underlying direct and inverse comorbidity. The figure summarises the study design and main findings. The central panel illustrates the concept of shared genetic architecture between neuropsychiatric disorders and cancers, manifesting as either direct comorbidity (positive genetic correlation, *r_g_ >* 0) or inverse comorbidity (negative genetic correlation, *r_g_ <* 0). Selected disease-pair examples are shown on the right. The lower panels detail the biological programs identified through local genetic covariance and pathway enrichment analyses: direct comorbidities are consistent with shared inflammatory, survival, tissue-remodelling and telomere-maintenance programs that may promote adaptation in both neuronal and tumour contexts, whereas inverse comorbidities suggest opposing selective pressures on apoptotic regulation, stress-response proteostasis, neuroprotection and immune-tone calibration between neuronal and tumour-cell states. The balance-scale motif represents the context-dependent equilibrium of these shared programs. The bottom panel summarises the contribution and translational relevance of the findings.

## 3 Discussion

This study provides a genome-scale genetic atlas of direct and inverse neuropsychiatric–cancer comorbidity, contributing a population-level inherited perspective to the emerging field of cancer neuroscience. By integrating global and local genetic correlation analyses with subtype-resolved and multi-omic comparisons across 35 diseases and 115 GWAS datasets, we show that the shared inherited architecture linking the nervous system and cancer is highly context-dependent and often obscured by broad diagnostic labels. Genomic data capture inherited liability upstream of downstream molecular states, making them useful for identifying stable cross-disease architecture before disease onset, treatment exposure or tissue degeneration distort the signal^23, 24^.

The predominance of positive genetic correlations and their concentration within disease categories is consistent with well-established transdiagnostic overlap across psychiatric diseases^24^. The positive overlap of psychiatric conditions with some cancers is plausible in view of the repeated implication of oxidative stress, inflammatory activation, altered neuroendocrine signalling and sleep disruption in psychiatric illness^73, 74^, all of which can be exacerbated through poor health behaviours and reduced self-care associated with neuroinflammation, tumour-promoting stress states or both^75, 76^. Adding a further mechanistic layer, neurotransmitters and neuropeptides dysregulated in psychiatric illness, including norepinephrine, dopamine and serotonin, can directly shape tumour-associated immune cells in the microenvironment, promoting myeloid-derived suppressor-cell recruitment and T-cell exhaustion^61^. By contrast, the most striking negative burden fell on cancer–neurological and neurodegenerative pairs, supporting the idea that inverse comorbidity between cancer and CNS disorders reflects a substantial biological component enriched in pathways involved in cellular proliferation, survival, stress tolerance and cell death, rather than only diagnostic or survival bias^77–79^. Consistent with this view, recent work has proposed that chronic neuronal stress and altered intercellular communication in neurodegeneration may reshape the tumour microenvironment through the neuro–immune–cancer axis, creating contexts that are less permissive to tumour growth^80^.

The local correlation analyses provide a mechanistic layer to these observations. The recurrence of specific loci across multiple disease pairs, notably chr9:126.9–128.9 Mb across ADHD–lung cancer and schizophrenia–lung cancer, chr2:103.2–104.5 Mb across ADHD–lung cancer and bipolar disorder–epilepsy through POU3F3, and calcium-channel loci in bipolar disorder–epilepsy, points to genomic modules that contribute transdiagnostically across psychiatric, neurodevelopmental and cancer-related phenotypes. Telomere maintenance, interferon signalling, vesicle trafficking, membrane excitability and FGFR-related pathways were particularly recurrent. FGFR signalling is also implicated in cellular migration and tissue patterning during nervous-system development and in invasive or metastatic cancer programmes, although these roles should be interpreted here as pathway-level hypotheses rather than direct evidence of cancer progression in the analysed cohorts**^?^ ^?^**. For direct comorbidities, these modules are consistent with shared survival programs, inflammatory signalling, cellular migration and tissue-remodelling pathways that may serve adaptive functions in both neuronal and tumour contexts. For inverse comorbidities, the data suggest that the same broad biological systems are tuned toward opposite optimal states: neuronal integrity and tumour fitness may impose conflicting demands on apoptotic regulation, immune tone and stress-response calibration, such that alleles favouring one state are in tension with the other. This model accommodates both the negative autism–prostate cancer signal concentrated in extrinsic apoptotic signalling (CASP8, CFLAR) and the ADHD–melanoma inverse correlation driven by opposing immune and UV-response architecture.

Several lung-cancer-positive axes identified here overlap with pathways already therapeutically actionable in non-small-cell lung cancer, including ALK, FGFR1, JAK1/interferon-related signalling and IDO1-mediated immune escape^58–60, 81^, suggesting that the shared genetic architecture is concentrated in pathways where pharmacological modulation already exists or is actively being developed, and that neuropsychiatric genetic data may offer an orthogonal route to target prioritisation in precision oncology. A full discussion of potential therapeutic prioritisation is provided in Supplementary Note 4.

Meta-analysis was particularly valuable because it recovered neuropsychiatric–cancer associations not visible in single-dataset scans, including Alzheimer’s disease with breast cancer and melanoma, Parkinson’s disease with breast and prostate cancer and ALS with schizophrenia, suggesting that some disease relationships are distributed across multiple datasets and modified by subtype, ascertainment or population composition. The negative Alzheimer’s disease–breast cancer estimates observed here contrast with an earlier cross-trait LD Score regression study, co-authored by Driver, that reported positive genetic correlations between Alzheimer’s disease and breast cancer and lung cancer using IGAP and GAME-ON summary statistics**^?^**. This discrepancy is biologically and methodologically informative, as it may reflect differences in GWAS releases, cancer subtype definitions, power, sample overlap, ancestry filtering or the meta-analytic aggregation strategy used here. The appearance of negative Alzheimer’s disease–breast cancer and positive Alzheimer’s disease–cervical cancer signals after aggregation is also consistent with recent bidirectional Mendelian randomisation work suggesting that the relationship between Alzheimer’s disease and gynaecologic cancers differs by tumour subtype^82^.

Perhaps the most consequential finding for the design of future neuropsychiatric–cancer genetic studies is the subtype divergence of familial and late-onset Alzheimer’s disease with respect to lung cancer and ADHD. Familial Alzheimer’s disease is dominated by penetrant amyloid-processing biology involving APP, PSEN1 and PSEN2, whereas late-onset Alzheimer’s disease is shaped more strongly by APOE, lipid handling, vascular ageing and immune regulation^42, 43^. These aetiological differences translate into opposing genetic correlations with cancer phenotypes and have direct practical implications. This pattern is independently supported by a recent systematic review of Mendelian randomisation studies, which found that the inverse association between Alzheimer’s disease and cancer is most consistent for breast cancer, especially oestrogen receptor-positive breast cancer, whereas evidence for other tumour types remains heterogeneous^35^. Genetic comorbidity studies that aggregate Alzheimer’s disease subtypes without distinction are therefore likely to obscure or cancel opposing neuro-oncological signals. The same caveat applies to any disease for which clinically or aetiologically distinct subtypes have been collapsed under a single phenotype label, and argues for subtype resolution as a methodological standard in future genetic atlases of neuropsychiatric–cancer comorbidity.

This biological heterogeneity is independently supported by the multi-omic convergence analysis. Benchmarking our results against epidemiologically established comorbidities from population-scale electronic health records^83^ shows that LDSC and HDL are competitive discovery tools, recovering a comparable or greater number of known comorbid pairs than transcriptomic, symptom or gene-based approaches despite operating on a fundamentally different layer of biological information. The convergence between genetic and transcriptomic evidence is particularly notable, as inherited susceptibility and downstream gene expression are sensitive to different confounding sources, and their agreement across the same disease pairs strengthens the case for a shared molecular aetiology. The pattern of convergence mirrors that of the genetic correlations themselves: psychiatric pairs are consistently recovered across multiple layers, reflecting both genuine biological overlap and the relative data maturity of these conditions, whereas inter-category pairs involving neurological or cancer diseases show lower multi-omic support and higher rates of missing data. Pairs recovered exclusively by genetic correlation but not by other layers may point to inherited comorbidities not yet visible at the molecular level, either because the relevant tissues or disease stages have not been adequately profiled, or because the shared architecture operates through regulatory rather than coding variation.

Taken together, our results argue that neuropsychiatric–cancer disease pairs should not be treated as fixed biological entities: subtype, sex, smoking exposure and family-history structure can substantially reshape the shared genetic architecture, and the direction of a global genetic correlation may conceal opposing local signals that only become visible when subtypes are considered separately. Integrating the genome-wide and local correlation approaches described here with transcriptomic, epigenomic, proteomic and epidemiological data, and doing so at the level of disease subtypes rather than aggregated phenotypes, represents the most tractable path toward resolving the subtype-specific mechanisms and population-specific risk profiles that underlie the neuropsychiatric–cancer comorbidity landscape described in this study.

### 3.1 Limitations

This study has several limitations. All analyses were restricted to GWAS datasets of predominantly European ancestry, which reduces heterogeneity but limits generalisability to populations with different allele frequencies, linkage disequilibrium structures and environmental exposures. In addition, the division of disorders into psychiatric, neurological and cancer categories was necessarily pragmatic rather than biologically absolute (Supplementary Note 1). Methodological constraints should also be considered. Although significance correction was applied against the full set of 6,555 possible trait pairs, LDSC and HDL used different reference panels, following individual method recommendations, and not all pairs could be compared owing to missingness. In a number of cases, inspection of the HDL output indicated a mismatch with the reference panel, causing estimates to collapse towards zero and likely explaining some isolated sign reversals. Likewise, several summary-statistic fields were reconstructed during preprocessing to harmonise heterogeneous GWAS releases. The transformation from odds ratio to *β* was empirically checked and did not materially alter results.

Further limitations arise from the study design. The meta-analysis combined genetic correlation estimates rather than the underlying GWAS summary statistics, and the evidential depth was uneven across diseases, with some phenotypes represented by only a single dataset and others by multiple datasets from a limited number of studies and partially overlapping cohorts. Although robust modelling reduces this dependence, residual non-independence may still affect precision. Finally, the local correlation and pathway enrichment results should be viewed as hypothesis-generating because they depend on the genomic partitioning framework, linkage disequilibrium reference and annotation strategy, and some recurrent loci still lack independent external validation. More broadly, genetic correlation captures only the component of comorbidity attributable to shared common-variant architecture and does not directly model rare variation, treatment effects, survival bias or temporally ordered disease trajectories.

## 4 Conclusion

Our study provides a genome-scale genetic atlas of direct and inverse neuropsychiatric–cancer comorbidity, offering a population-level inherited perspective on the biological programs that may link the nervous system and cancer. By integrating global and local genetic correlation analyses with subtype-resolved and multi-omic comparisons, we show that shared disease architecture is highly context-dependent and often obscured by broad diagnostic labels. Psychiatric disorders displayed the strongest and most reproducible genetic overlap, whereas several cancer–neurological associations pointed to inverse relationships consistent with competing demands on apoptotic control, immune tone and cellular stress responses, biological tensions that have remained poorly characterised at the genome-wide level.

Beyond defining broad patterns of convergence and divergence, these analyses identify recurrent genomic loci and inherited biological programs that may underlie disease co-occurrence or mutual exclusion between neuropsychiatric and cancer phenotypes. The subtype-dependent architecture revealed here, most strikingly in the opposing neuro-oncological profiles of familial and late-onset Alzheimer’s disease, has direct implications for the design of genetic comorbidity studies and argues for subtype resolution as a methodological standard in future research on neuropsychiatric–cancer relationships. Several of the shared pathways identified, including interferon–JAK–STAT signalling, FGFR cascades and apoptotic regulators such as CASP8 and CFLAR, are already therapeutically actionable in cancer, suggesting that inherited neuropsychiatric–cancer genetic architecture may offer an orthogonal route to target prioritisation in precision oncology.

These findings establish that the genetic basis of neuropsychiatric–cancer comorbidity is neither uniform nor reducible to shared risk factors, but reflects a structured landscape of inherited biological programs whose direction and magnitude depend on disease subtype, sex and ascertainment context. We anticipate that integrating this genome-scale map with mechanistic, epigenomic and single-cell data will accelerate the identification of the molecular nodes at which the nervous system and cancer converge, and diverge, in ways that are both biologically fundamental and therapeutically exploitable.

## 5 Methods

### 5.1 Data collection, preprocessing and heritability estimation

Genomic data were downloaded from the GWAS Catalog and the Psychiatric Genomics Consortium, focusing on datasets describing a single condition within three major groups: cancer, psychiatric and neurological or neurodegenerative disorders.

To minimise variability bias and confounding, we restricted the selection to individuals of European ancestry, the most prevalent ancestry in the catalogue (approximately 91%)^84^. Only genome-wide genotyping-array studies with at least 4,500 cases and a total sample size of 10,000 individuals were selected. Datasets were also required to include a reference SNP cluster ID and columns with summary statistics for effect size and significance, including *z* score, *P* value, *β* and odds ratio.

We compiled 123 GWAS summary-statistic datasets spanning 35 unique diseases, including 112 from the GWAS Catalog and 11 from the Psychiatric Genomics Consortium. To ensure consistency across all datasets, we developed a preprocessing pipeline to harmonise GWAS summary statistics (Supplementary Note 5 and Supplementary Table 5). Missing *β* values were derived from odds ratios, *z* scores from *β* and *P* values, and standard errors of *β* from odds-ratio confidence intervals. Datasets were subsequently harmonised into the specific formats required by LDSC, SUPERGNOVA and HDL. Following LDSC developers’ recommendations, heritability *z* scores were calculated for each trait, and datasets with a *z* score below four were removed to avoid noise from low heritability^85^, resulting in the removal of 8 datasets and a final total of 115. Each dataset was assigned to a selection tier: main (largest sample size per disease; *n* = 34), minor (additional non-subtype datasets; *n* = 40), subtype (clinically defined subtypes; *n* = 30) and subtype-family (familial case subsets; *n* = 11). Diseases were grouped into three broad categories: cancer, neurological or neurodegenerative and psychiatric (Supplementary Fig. 7, Supplementary Table 6 and Supplementary Note 1).

### 5.2 Global genetic correlation inference

To estimate the genetic correlation (*r_g_*) between disease pairs, we used two complementary approaches: LDSC^85^ and HDL^86^. LDSC quantifies the extent to which SNP effect sizes are correlated across traits while accounting for linkage disequilibrium patterns, thereby distinguishing true polygenic signal from confounding due to population stratification. HDL leverages linkage disequilibrium information from a large reference panel to model the covariance structure of test statistics genome-wide, increasing statistical power and reducing bias, particularly for highly polygenic traits^87^. Using both methods in parallel allowed us to balance LDSC’s robustness with HDL’s enhanced sensitivity. Multiple testing was corrected using the Benjamini–Yekutieli approach^88^. The ability to obtain significant correlations across the three disease categories was compared using Fisher exact tests. Jaccard indices were calculated to assess agreement between LDSC and HDL results at the global and disease-by-disease level.

### 5.3 Meta-analysis of global genetic correlations

We meta-analysed genetic correlation estimates for each disease pair using the metafor R package^89^. Analyses were performed without subtypes, which were used only for downstream interpretation. When multiple estimates were available, we preferentially fitted a multivariate random-effects model with *t* test inference, requiring at least five estimates and two unique clusters per identifier. Otherwise, a univariate random-effects model with Knapp–Hartung adjustment was fitted. Cluster-robust standard errors were computed using clubSandwich CR2 and, when necessary, metafor::robust.

### 5.4 Local genetic covariance and correlation inference

Local genetic covariance and correlations were calculated using SUPERGNOVA^90^, which partitions the genome into independent loci and applies GWAS summary statistics to estimate region-specific pleiotropy. Regions with *P ≤* 0.05 were considered significant. Gene annotation and enrichment analyses were performed using rGREAT^91^, applying region-based functional enrichment with hg19 transcription start site definitions against Gene Ontology Biological Process^92^ and Reactome pathways^93^.

### 5.5 Multiomic comorbidity recovery analysis

To evaluate whether the genetic correlations identified in this study converge with evidence from other biological layers, we assembled a multiomic comorbidity matrix for the disease pairs included in our dataset. As a reference set of established comorbidities, we used disease pairs identified from Danish electronic health records^83^ that overlapped with the diseases analysed here. Disease names were mapped to ICD-10 codes to enable harmonisation across all data sources. Recovery was assessed only for positive comorbidities with relative risk greater than 1.01.

For each disease pair, we collected evidence of similarity or co-association from seven independent omics layers and two gene-based layers derived from GWAS data. For all molecular layers, pairwise disease similarities were computed using a common weighting framework: each molecular entity (gene, drug, miRNA, metabolite, microbe or symptom) was assigned a frequency score, defined as the number of diseases associated with that entity divided by the maximum number of diseases associated with any entity in the dataset. Similarities between all disease pairs were then computed using either weighted Jaccard indices or cosine similarities, depending on the layer, with statistical significance assessed via randomisation (*n* = 10, 000 permutations) and multiple-testing correction using the Bonferroni approach. Pairs with FDR greater than 0.05 were set to zero before network construction. The specific metric used for each layer was determined by the file loaded in the analysis pipeline.

GWAS gene–disease associations were derived from the GWAS Catalog by mapping significantly associated loci to genes for each disease in our dataset. Pairwise disease overlap was quantified as the raw intersection of mapped genes between each disease pair. DisGeNET disease–gene associations were extracted from manually curated associations^94^, and pairwise disease overlap was quantified as the raw intersection of associated genes. Disease–drug associations were extracted from ChEMBL, retaining only curated therapeutic associations and excluding inferred associations^95^. Disease–miRNA associations were extracted from HMDD, retaining only manually curated associations from the literature^96^. Disease–metabolite associations were extracted from the Human Metabolome Database, retaining only blood metabolite measurements^97^. Disease–microbe associations were extracted from DISBIOME, retaining only gut microbiome associations^98^. Disease–symptom co-occurrence data were generated following the approach of Zhou et al.^99^, extracting co-occurrences between diseases and symptoms from PubMed metadata updated to March 2025. Significant disease–symptom associations were identified by the chi-square test. Transcriptomic inter-disease similarity scores were obtained from Sánchez-Valle et al.^20^, based on pairwise cosine similarities between diseases computed from differential gene-expression fold-change vectors. LDSC and HDL genetic correlation results from the present study were included as two additional evidence layers, using FDR-corrected significant positive pairs as the recovery criterion. For each omic source, a disease pair was classified as recovered if a statistically significant positive similarity was detected, as undetectable if at least one disease in the pair lacked data in the corresponding resource and as not recovered if data were available for both diseases but no significant similarity was identified. The total number of layers recovered for each pair was summed to produce a convergence score ranging from 0 to 10.

## Author contributions

M.F.-R. and J.S.-V. designed the study and performed the analyses. R.T.-S., A.V., S.M., F.W., J.F.-M. and J.V.-S.O. provided technical advice. M.F.-R. and J.S.-V. wrote the manuscript. All authors discussed the results and commented on the manuscript.

## Data availability

All data needed to understand and assess the conclusions of this research are available in the main text, Supplementary Information and the SOROLLA atlas at http://147.156.4.159:3838/sorollaatlas/. Raw GWAS summary statistics are publicly available from the GWAS Catalog and the Psychiatric Genomics Consortium.

## Code availability

Analysis code is available at https://github.com/MFloresRodero/SOROLLA-gencor.git.

